# Curvature coding in early visual system revealed by scale variance during adaptation to flashing circles

**DOI:** 10.1101/2023.02.11.528121

**Authors:** Irfa Nisar, James H. Elder

## Abstract

How is curvature coded in the early human visual system? Humans are successful in recognizing objects and by extension, the shape representing the object, under varying scale conditions (Biederman & Cooper, 1992; Lindeberg, 2013). How do we neuro-physiologically code the invariance (or variance) in curvature and does the curvature coding change with scale? The circle-polygon illusion produces polygonal percepts during adaptation when a static dark outline circle is pulsed at 2 Hz alternating with a gradient luminance circle. We use the circle-polygon to study curvature processing with respect to size and scale. Both the radius and eccentricity of the stimulus were varied in a crossed design over 1-8 deg. Observers reported a circle or the polygon order and the strength of the percept. We test a lower level account that argues for curvature opponency between neurons against a higher level account that codes for whole shapes. This higher level account supports scale invariance, a property through which we recognize objects regardless of the object’s size on the retina. We show the following: (1) Scale invariance is not obeyed during adaptation. The mean order of the perceived polygon increased with stimulus size and decreased with eccentricity. This also demonstrates that curvature coding occurs in the early visual system. (2) Linear regression analysis reveals that the cortical size of the stimulus is a better predictor of perceived polygon order. We quantify the relationship parametrically between cortical size and polygon order. Using integration and regression, we identify the region of the cortex, V1, where the shape, a regular ordered polygon, is being computationally constructed.

## Introduction

Why is curvature perception important? Coding of curvature through the visual system influences how complex shapes are recognized in the world and why the human visual system is able to identify the same shape presented at varying scales (Biederman & Cooper, 1992; Lindeberg, 2013). Rotation invariance is observed in region V4 of the visual cortex with respect to normalized curvature (El-Shamayleh & Pasupathy, 2016a). Size and rotation invariance is modelled by rotationally symmetric and linearly separable Gaussians and its partial derivatives as a representation of the lower visual hierarchy, LGN and V1 receptive fields (Lindeberg, 2013). The research objective is to answer the following questions that probe curvature processing with respect to scale: (a) Is curvature coded on a low level retinal plane or is it a product of a higher order cortical process? (b) Is the coding stimuli-scale variant or invariant: Does the size of the stimuli change the number of sides of the polygon seen in the percept after adaptation? (c) Does the polygon order (shape) change with size and eccentricity? Can we quantify the relationship between the shape seen during in-place adaptation and the independent variables, size and eccentricity, at which the stimuli is presented? (d) Is the polygon orientation rotationally invariant? We associate the canonical position associated with a polygon as the position when the polygon has a vertex at the 12 o’clock position. As shown in this paper, users do not show a pronounced preference for the canonical position or for a particular orientation away from the canonical position.

### Circle Polygon Illusion

We adopt the circle-polygon illusion that was discovered by Khuu (Khuu, 2002) and use Sakurai’s technique (Sakurai, 2014; Sakurai & Beaudot, 2015) to construct the phenomenon to understand how curvature is coded in the brain. Sakurai’s illusion, builds on Khuu’s circle-appearing-as-polygon illusion, by temporally pulsing a graded circle with its outline at 2 Hz. The circle-polygon illusion demonstrates that polygons are produced after prolonged adaptation to flashing circles. Although (Khuu, 2002; Sakurai, 2014; Sakurai & Beaudot, 2015) only report hexagon as the emergent shape, Ito in (Ito, 2012) presents an array of shapes (triangle,square,pentagon,hexagon,heptagon and circle) as the independent variable response type with several responses that are not hexagons. (Ito, 2012) shows that adaptation to a static and a rotating circle,either filled or unfilled, generates an afterimage of a hexagon. In (Sakurai, 2014; Sakurai & Beaudot, 2015), Sakurai show that if a circle and its inward gradation are alternated, we see a polygon shape perception of the circular stimuli itself. That shape is a hexagon.

### Lower level or higher level?

The early visual system is defined by the retina, and controlled through the receptive fields of regions V1 through V2 (D. H. Hubel & Wiesel, 1959; D. H. Hubel, Wiesel, & Stryker, 1978; Hegdé & Van Essen, 2000). The higher mid-level V4 neurons, with larger receptive fields, while showing preference for certain angles, show scale-invariance to different sized arc boundaries that subtend the same angle at the stimulus’s center (Pasupathy & Connor, 2001, 2002; El-Shamayleh & Pasupathy, 2016a). We probe systematically across eccentricity and size to see if this scaling phenomenon holds and if not, can we deduce a relationship that explains the scaling. Ito (Ito, 2012) assert that the circle-polygon phenomenon is cortical because interocular transfers result in users reporting circles whereas each eye under monoptical conditions, when shown the same circular stimuli would produce a polygon. Sakurai in (Sakurai & Beaudot, 2015), build on Khuu’s circle-appearing-as-polygon illusion and also cite a cortical process is responsible given that the adaptation occurs almost instantly when the stimuli pulsation is varied between 0.5 Hz to 8 Hz. Beyond the observation that it is cortical, can we quantitatively state that polygons produced are determinant of the size and eccentricity as a known function of a certain region of the cortex? This would explain curvature processing for future shape description without undergoing adaptation. Several regions of the visual cortex show invariance to scale and selective invariance to rotation. If the polygonal response to curvature was coded on the retinal plane or lower levels V1 and V2, it would be influenced by the neuronal responses to the shape as it appears on the retina. Size and rotational invariance is modelled by rotationally symmetric and linearly separable Gaussians and its partial derivatives (Lindeberg, 2013) as a representation of the lower visual hierarchy receptive fields found in the LGN and V1 regions. If it was coded higher up in the cortex, such as the mid level V4, it might show more complex representations such as scale invariance with respect to normalized curvature (Pasupathy & Connor, 2001, 2002; El-Shamayleh & Pasupathy, 2016a). Size invariance is also a function of the higher visual hierarchy, beyond V4, observed at the inferotemporal (IT) region (Biederman & Cooper, 1992). Schwartz et. al (Schwartz, Desimone, Albright, & Gross, 1983) demonstrate invariance by recording from the inferotemporal (IT) region, a region that codes for global shape, by showing that position and size invariance is observed for stimuli constructed from Fourier descriptors.

### Regularization or Regular polygons

As seen in (Sakurai, 2014; Sakurai & Beaudot, 2015), the circular stimuli, when alternated with its inward gradation, makes the cortical process adapt to the curvature and gives rise to symmetrical regular hexagons instead of an irregular organic shape. Gheorghiu and Kingdom in (Gheorghiu & Kingdom, 2007, 2009, 2017) demonstrate that population neural code affects perception of local curvature through an AND-gate multiplication mechanism. However, none of the above experiments explain the order of the polygon as a function of the stimuli size. Neurophysiological studies conducted on non-primate data in (Das & Gilbert, 1995) show that receptive fields that fire for a similar range of orientations cluster together but long-range connections can sometimes influence the range of degree to which orientations are recognized. Why is the regularity of the afterimage, also called regular polygonalization, an important attribute in the afterimage? The presence of regularization itself suggests that retinal bleaching or retinal adaptation (Spillmann, 1994; Schrauf, Lingelbach, & Wist, 1997) is an unfavourable hypothesis since bleaching produces disappearances in features or production of new ones (opposite coloured fault lines) when intensity rods are fatigued (Spillmann, 1994; Schrauf et al., 1997). The disappearance occurs peripheral to fixation where fatiguing is pronounced and lacks the deliberate production of symmetry which could only be produced if rods or neurons were aligned in a corresponding symmetrical pattern. We test if a cortical process operating in the lower regions such as V1 utilizes long range connections in the production of polygonal shapes. In this paper, we compare a retinal and a competing cortical model to see which better explains the adapted polygon and if we can quantitatively define a relationship. The retinal model relates the retinal size of the stimulus to the order of the polygon. The second model, relies on cortical M-scaling and relates the cortical image of the stimulus to the order of the polygon.

The visual cortex is a large area with several sub-regions V1, V2, V3 and V4. We use behavioural criterion as the dependent variable and neuroscience theory to construct what the stimulus’ circumference looks like on the visual cortex and which region best explains the edge length being reported for each stimulus.

### Parametric cortical equation

How can one quantify the stimuli’s projection on the cortex? It is known that receptive field size increase towards the retina’s periphery (Rovamo & Virsu, 1979). M is defined as the ratio between the cortical representation of the receptive field (in mm) and its size on the retina (degrees of visual angle). In (Daniel & Whitteridge, 1961), Daniel and Whitteridge show that an increment of 1 degree visual angle measured from the central fixation point corresponds to an arc length of M mm on the theoretical visual cortex.

Rovamo and Virsu in (Rovamo & Virsu, 1979) estimate M, for the principal meridians of the visual field in equation 1 and the accompanying table for coefficients. Here, *M*_*o*_ is 7.99*mm/*^°^, the magnification at the fovea. *r* is the eccentricity in degrees.

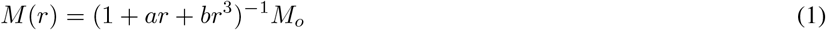

| Area where magnification is computed | a | b | Eccentricity range |
| --- | --- | --- | --- |
| Nasal | 0.33 | 0.00007 | $0 \leq E \leq 60 \text{ deg}$ |

If we integrate M, we obtain the distance on the cortex as a function of eccentricity. Our skeletal algorithm, see Algorithm 1, computes the integral on the cortex and is explained in the Methods section.

## Experiment

This section is divided in two parts. We first describe the theory to compute the shape on the cortex using M-scaling. This requires behavioural responses that elicit the polygon seen during in-place adaptation. Next, we collect the behavioural responses that answer questions about scale, orientation and shape during in-place adaptation. The experiment reports the polygon percept seen during adaptation as the circle radius and the eccentricity of the circle’s placement varies.

## Methods

### Computational Overview

Neuroscience experiments do not describe shapes as it would appear on the cortex after M scaling. Most concern themselves with studying activation areas given a suitable stimuli (Sereno et al., 1995; Harvey & Dumoulin, 2011) or mapping out the biological receptive fields that would support a change of orientation (thereby describing curves) across cortical columns (Das & Gilbert, 1997). As such, only localized or generalized description of possible shape outcomes on the cortex exist. In this paper, we attempt a description of cortical shape by using cortical M-scaling.

In order to explain if the polygons observed above can be explained quantitatively by a retinal model or a higher level cortical model, we apply linear regression. We fit a first order linear model of the polygon order as a function of retinal size and another as a function of cortical circumference size. A retinal function samples the circular stimulus in 2D cartesian co-ordinates. The domain of the function is degree visual angle representing eccentricity. The cortical length can be computed from the points on the circle seen on the 2D retinal plane and applying the cortical M-scaling, that measures millimetres of cortex per degree of visual angle, as a function of eccentricity, on those sampled 2D points.

We shall hereby call the retinal circumference size, retinal length, and the cortical circumference size, cortical length. See Figure 11 (a,b,d).

The M-scaling factors in the cortical function were borrowed from (Rovamo & Virsu, 1979). This is a skeletal model with Rovamo and Virsu cortical magnification factors. The cortical magnification factors can be replaced by other magnification factors that perform hierarchical sampling, such as when the cortical representation in V4 is represented by how the region in V1 is sampled (Harvey & Dumoulin, 2011). Harvey and Dumoulin create a model that maps population receptive fields and cortical magnification factor as a function of the cortical magnification factor of the first field in the visual hierarchy, V1. Harvey and Dumoulin (Harvey & Dumoulin, 2011) show that the cortical magnification factor in area V4 is smaller than those in V2 and V3 which was in turn is less than V1. The cortical magnification factor for fields such as V2,V3 and V4 that sample from V1 is independently constructed. We, therefore, use V4’s cortical magnification factor when considering if polygon order reported could be a function of cortical scaling in region V4, which is known for curvature representation.

In Algorithm 1, we approximate the cortical length formed by sampling the retinal stimuli’s circumference at a fixed number of points and projecting that sampled length onto the cortex. The length formed on the cortex is scaled by the magnification factor, M, to give the cortical length.

#### Algorithm 1

Cortical mapping function using a neurophysiological derived cortical magnification factor Compute the cortical length using the M-scaling magnification factor for a circle of a given radius and presented at a particular eccentricity.

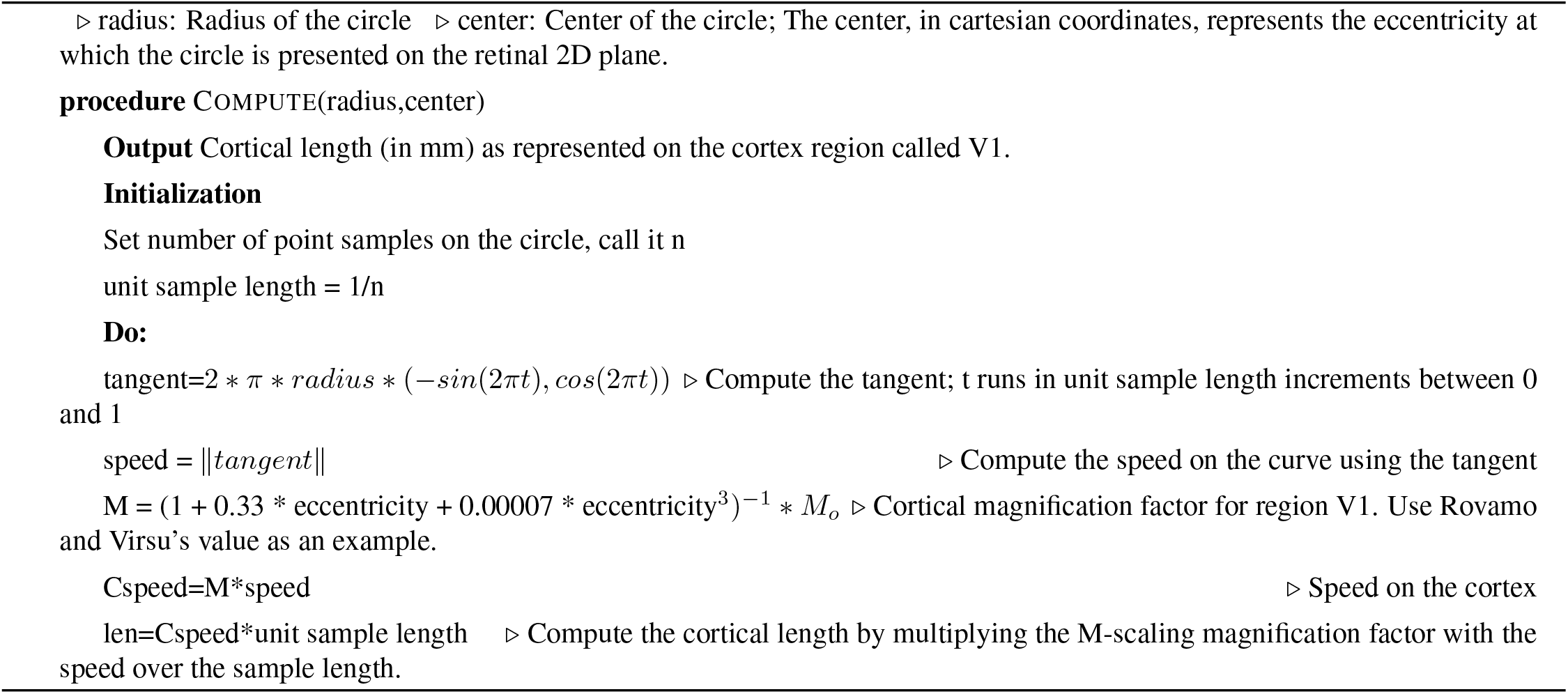

We use boot strapping and use the coefficient of determination, *R*^2^ to estimate the goodness of fit when fitting a linear regression function, polygon order as a function of the retinal and cortical length.

### Design

The main computational question centres around finding the region of the cortex that is responsible for the polygon being reported. In order to answer this question, we need the polygon order reported as a behavioural response. We need to verify if the eccentricity is a contributing effect in the polygon order formation in order to justify using M-scaling in the construction of shape. We use the polygon order and regress it against the circumference on the retina and the cortex. The circumference on the cortex is M-scaled such that it’s analytical projection is different in each of the region V1 and V4 based on the scale factor used.

We also try to understand the effect of size, eccentricity and participants on polygon order and strength, the two responses collected from participants. The experiments employ a within subjects design with polygon order as the dependent variable. We want to understand the effect of size, eccentricity and participants on the polygon order being reported. We use fitlme to model the data and use loglikelihood to address if the model fits the data well. We test significance of the effects using the p-value and the F-statistic reported. We want to know if strength and polygon order are co-related as strength could influence the perceptual formation of polygon order. We test this using Pearson’s correlation. In order to understand if the polygon reported had strength as a contributing effect, we let polygon order become the dependent variable and strength, the independent variable and test that model using fitlme. We test significance of the effects using the p-value and the F-statistic reported.

Lastly, we understand the effect of size, eccentricity and participants on the strength of the percept with strength the dependent variable and size and eccentricity the independent variables. We use fitlme to model the data.

To summarize the objective, as mentioned earlier, we want to answer the following questions that probe curvature processing with respect to scale: (a) Is curvature coded on a low level retinal plane or is it a product of a higher order cortical process? (b) Is the coding stimuli-scale variant or invariant: Does the size of the stimuli change the number of sides of the polygon seen in the percept after adaptation? (c) Does the polygon order (shape) change with size and eccentricity? Can we quantify the relationship between the shape seen during in-place adaptation and the independent variables, size and eccentricity, at which the stimuli is presented? (d) Is the polygon orientation rotationally invariant?

We used ANOVA for testing if the presentation of stimulus on the left or right of the visual field effected strength.

### Observers

Nine graduate students (6 male, 3 female) at the university participated in the experiment. Four of the graduate students had prior experience in participating in psychophysical experiments. The remaining five students had limited or no experience related to psychophysics. All the students had normal or corrected to normal vision. The experimental data and source code are stored at the Open Science Framework at the following link https://osf.io/x8n3p/.

### Stimuli

The stimuli consisted of a circle with a black outline that alternated with another circle of the same size but with an inward gradation pattern. The black outlines were constructed of penwidth 5 using Psychtoolbox. See 1 (b). The circles were built of different sizes. The diameter of the circle was 2,4,8 or 16 ^°^. See Figure 1 (a). The eccentricity at which the circle was presented was 1,2,4 and 8 ^°^. See Figure 2. Every unique combination of eccentricity and radius was presented ten times for a total of 160 trials. The trails were divided into 10 blocks with each block presenting each combination of size and eccentricity once. The presentation of the sixteen stimulus within a block was randomized. Users were allowed to continue to the next block when they were ready. The circle was placed left of the central fixation cross half the time and on the right of the fixation cross half of the time. The stimuli was presented against a white background.

**Figure 1:**
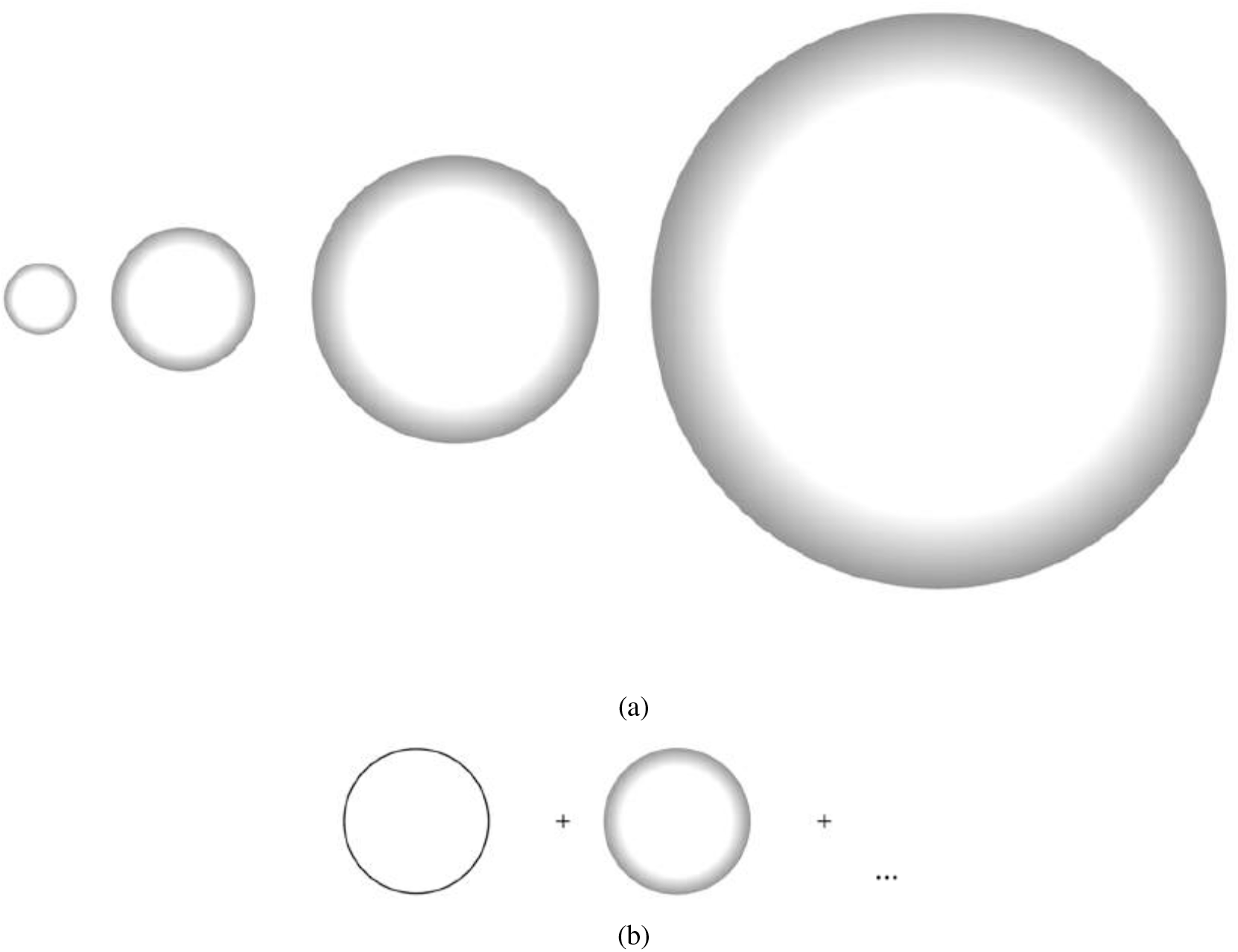
(a) Stimuli size: Circles,with inward gradation, whose size (diameter) subtends 2,4,8 and 16 degrees of visual angle respectively on the user’s eye. The pictures are for illustration only and show the relative sizes of the stimuli. Stimuli do not subtend the exact degree on the reader’s eye. (b) Flashing stimuli: Circle in a black outline and its inward luminance gradation. The outline and the inward gradation alternate at 2Hz. The circle is presented to the left of the fixation cross.

**Figure 2:**
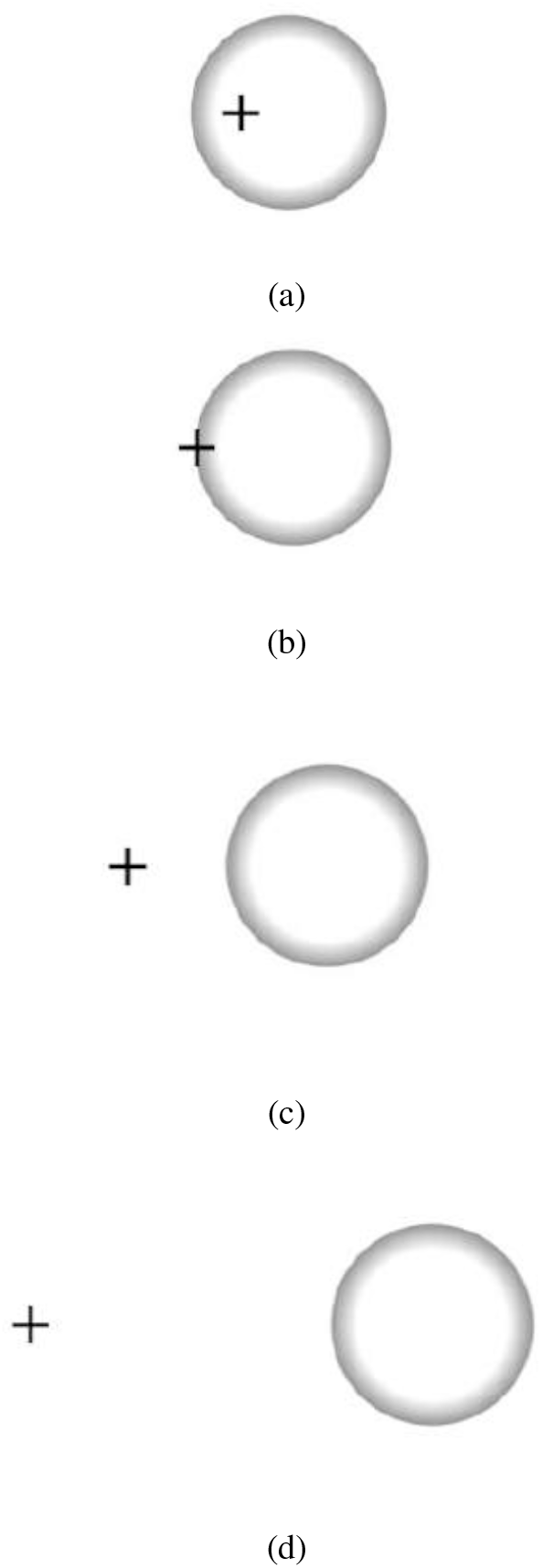
Presentation eccentricity: Circles,with inward gradation, presented at eccentricities of 1,2,4 and 8 degrees of visual angle respectively on the user’s eye. The circle’s size (diameter) is 4 degrees.

### Apparatus

Stimuli were generated using the Psychtoolbox (Brainard, 1997) in Matlab on a Windows machine with a Quadro K2200 graphics card and a DELL P2715Q LCD screen. The stimuli were presented on the same machine in the lab. A chin rest was used to ensure stable head position. No eye trackers were used. Instead the participants were instructed to fixate at the central cross without moving their eyes. The monitor resolution was 3840 pixels by 2160 pixels with a refresh rate of 60 Hz. The distance of the observer from the center of the monitor screen was 60 cm. The background was white. The mean luminance of the monitor was 26 cd/*m*^2^.

Psychtoolbox’s *Screen* function was used to create stimuli. The distance of the observer from the center of the screen and the size of the screen, both in centimeters, were used to compute the angle in degrees subtended at the user’s eye. Using the monitor’s resolution in pixels, the pixel per degree was computed. The *Screen* function was used to draw the appropriate stimuli using the pixel per degree information.

### Procedure

#### Measurement Techniques

The testing took place in a darkened room. Observers used a standard mouse with a middle mouse button and a left button. The participants indicated the polygon to which they adapted, the polygon’s orientation and its strength. Once the participant recorded a response, they pressed the middle mouse button on the mouse to move to the next stimuli presentation. The experiment was run in 5 blocks. After a block of 32 stimuli, a break was awarded to the observer. They could resume the next block of trial when ready. The left mouse button was used for all response recording tasks such as selecting the shape, its orientation and the strength of the percept (Figure 3).

**Figure 3:**
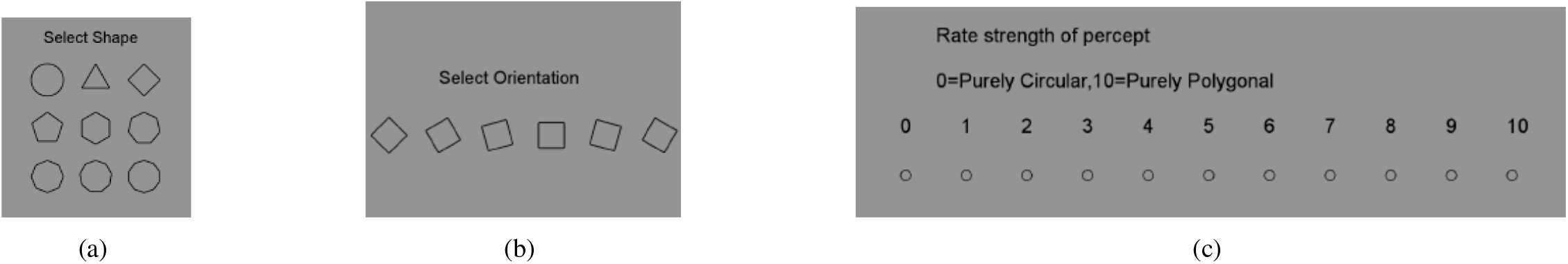
(a) Percept selection: Users select the in-place adaptation shape from this list of n-gons: triangle, square, pentagon, hexagon, heptagon, octagon, nonagon, decagon. (b) Orientation of the selected polygon: Users select the orientation of the polygon selected in increments of 15 deg. The above shows the orientations for a square. (c) Strength selection: The participants mark the strength of the percept on a scale from from 1-10 if they select a polygon. A circle is automatically entered as a zero and no likert scale is shown for a circular selection.

## Results

The results from the nine observers is shown in Figure 4. On average, most observers report more polygons than circles during the entire experiment. Polygons are reported 61.9 % of the time versus circles that are reported 38% of the time. This means that the formation of polygonal percepts is a phenomenon that occurs in the sampled population. A Student’s t-test confirmed no significant difference in tendency to perceive polygons between displays shown to the right or left of observer’s fixation (t(x) =T, p=.75). This rules out any effect of anisotropy in peripheral vision, asymmetric foveal confluence influencing shape formation and in the difference between the left and right visual field (Schwartz et al., 1983; Schira, Tyler, Breakspear, & Spehar, 2009). Anisotropy would distort the shape percept on the left and right. However, this test reveals that the emergent percept does not suffer from anisotropic shape distortions based on the strength of perception in the left and right visual fields. The strength is an effect for the polygon order reported (p *<* 0.05, t=29.54) in a test that employs fitlme with polygon order as the independent variable and strength as the dependent variable. A Pearson’s correlation (p *>* 0.05, r= 0.6144) shows that polygon order and strength are indeed correlated. However, this does not necessarily imply that lower strength gives rise to smaller order polygons or that the highest strength gives rise to the largest polygons as evident in Figure 4 (c).

**Figure 4:**
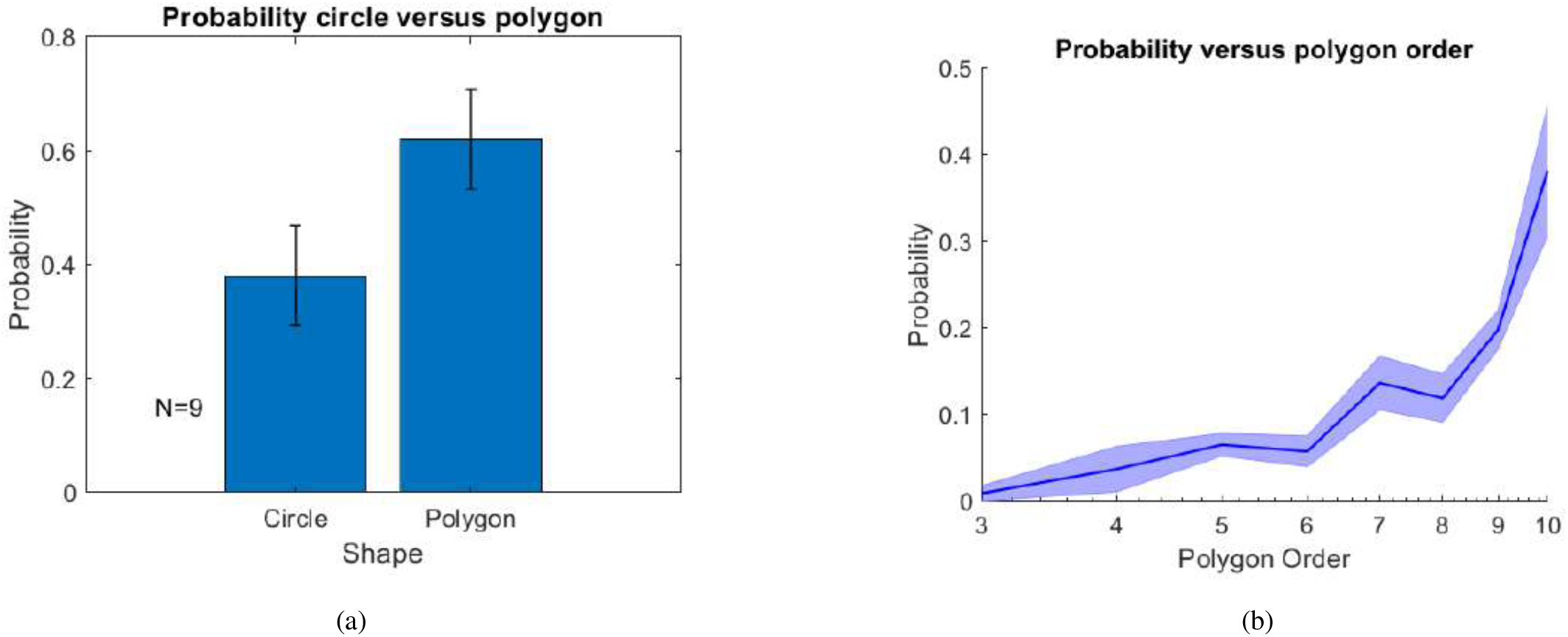
(a) Proportion Polygon: Observers report seeing polygons after adaptation 61.94 % of the time and seeing the same circle as the stimulus 38.06% of the time across all the stimuli combinations and trials presented. Error bars indicate the standard error of mean (SEM) around the mean proportion polygon (proportion polygon is reported per participant and averaged). (b) Probability distribution of the different polygon orders across all participants. Shaded error bars indicate the standard error of mean (SEM) around the mean polygon order.

### Strength

The experiment measured the strength of a percept on a scale from 0 to 10. 0 meant that the observer did not see a polygon percept after adaptation. The percept in this case remained the same as the circular stimuli. A 10 on the scale indicated a crisp regular polygon. Figure 5 (a) shows that observers do not report a difference between the left and right visual receptive fields when adapting to the stimuli. The strongest effect is for small objects at large eccentricities. See Figure 7 (a).

**Figure 5:**
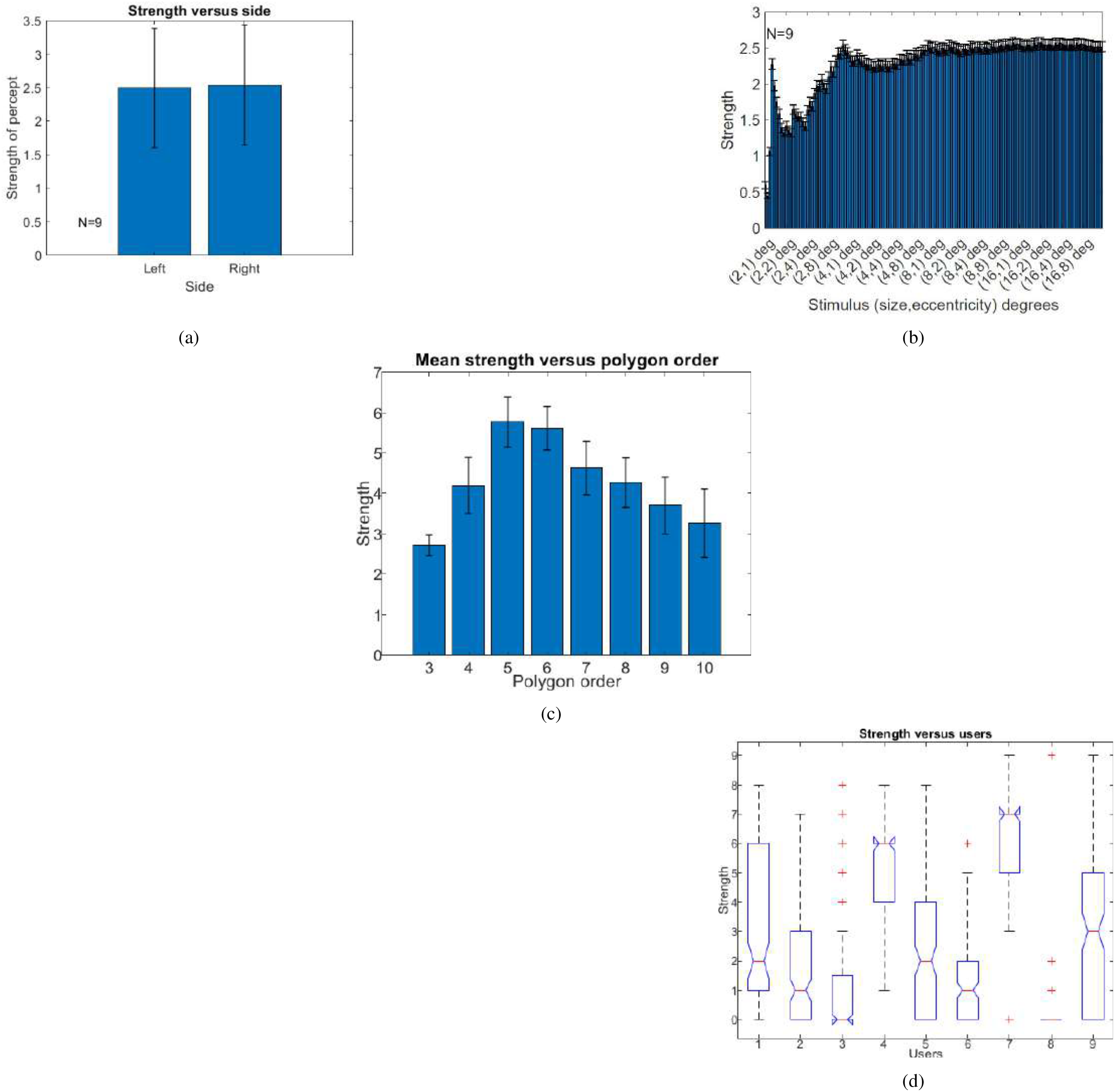
(a) Strength Versus Side: Users do not report any significant differences between the left and right visual receptive fields. The same stimuli is presented on the left and the right. Panel ‘Left’ depicts mean strength reported for stimulus presented to the left of the fixation cross. Panel ‘Right’ depicts mean strength reported for stimulus presented to the right of the fixation cross. Error bars represent standard error of mean (SEM) around the mean strength, reported across all participants. (b) The mean strength remains at 2.5 across almost all stimulus size and eccentric presentation. The only exception is observed with the smallest stimulus : size 2 deg that shows certain deviations across users. Users show a wide variation in the strength reported, indicating that there is a lack of uniform percept formation for the smallest stimulus sizes across users but becomes uniform as the stimulus becomes bigger. Error bars indicate the standard error of the mean centred around the mean. (c) Mean strength reported when polygon order indicated is selected as percept. Error bars indicate the standard error of mean around the mean. (d) Mean strength (Y-axis) reported by each participant (X-axis) across all trials of stimuli. The dash line represents the minimum and maximum strength reported. The bars represent standard deviation around the mean.

**Figure 6:**
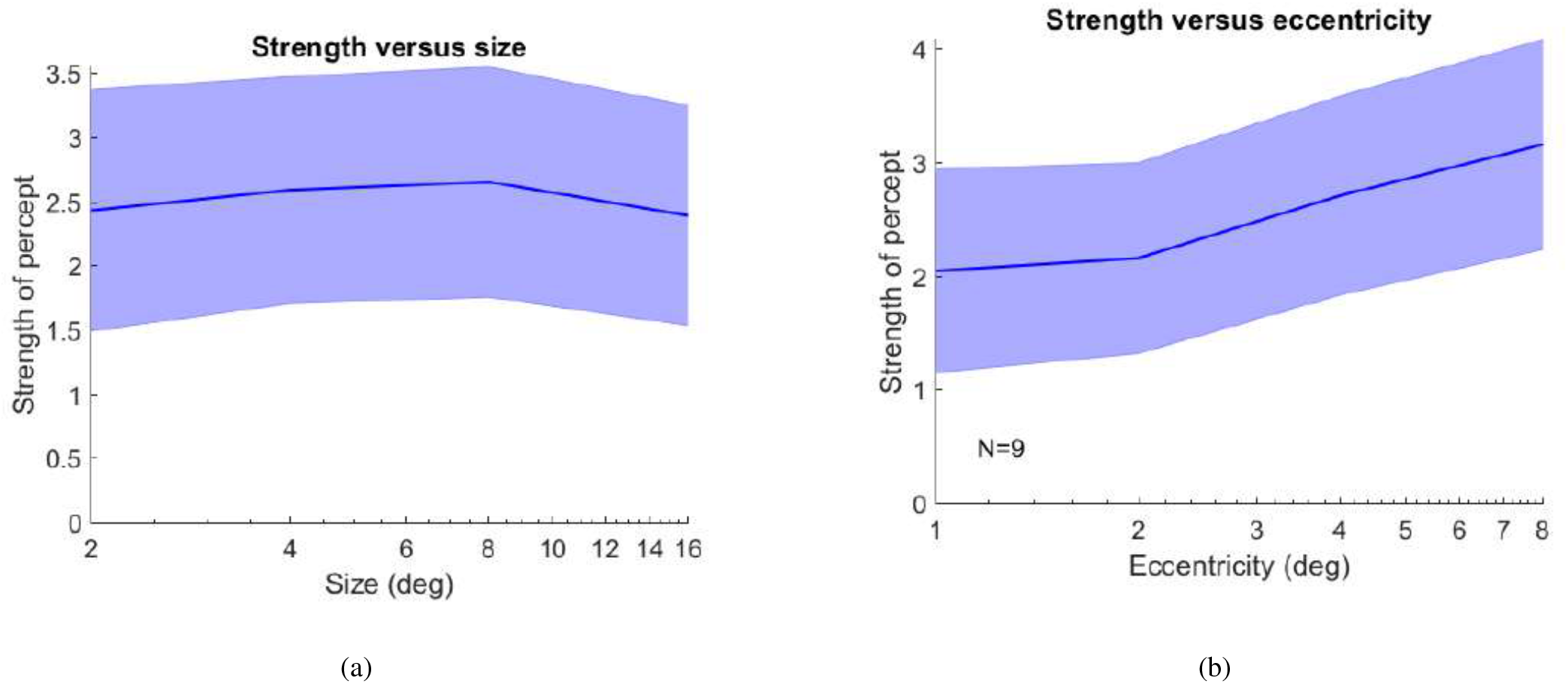
(a) Strength versus size: Mean strength reported by users when shown a stimulus of size(diameter) of (2,4,8,16) deg visual angle. Error bars represent standard error of mean (SEM) around the mean. Users, on average, report the sharpest perception of the polygon when the radius of the polygon subtends 4 degrees of visual angle on the user’s eye. (b) Strength versus eccentricity: Mean strength reported by users when stimulus is placed at an eccentricity of (1,2,4,8) deg visual angle. Error bars represent standard error of mean (SEM) around the mean. Users report the sharpest polygon responses when the stimuli is presented at the largest degree (8 deg) of visual angle. Error bars indicate the standard error of mean interval around the mean.

**Figure 7:**
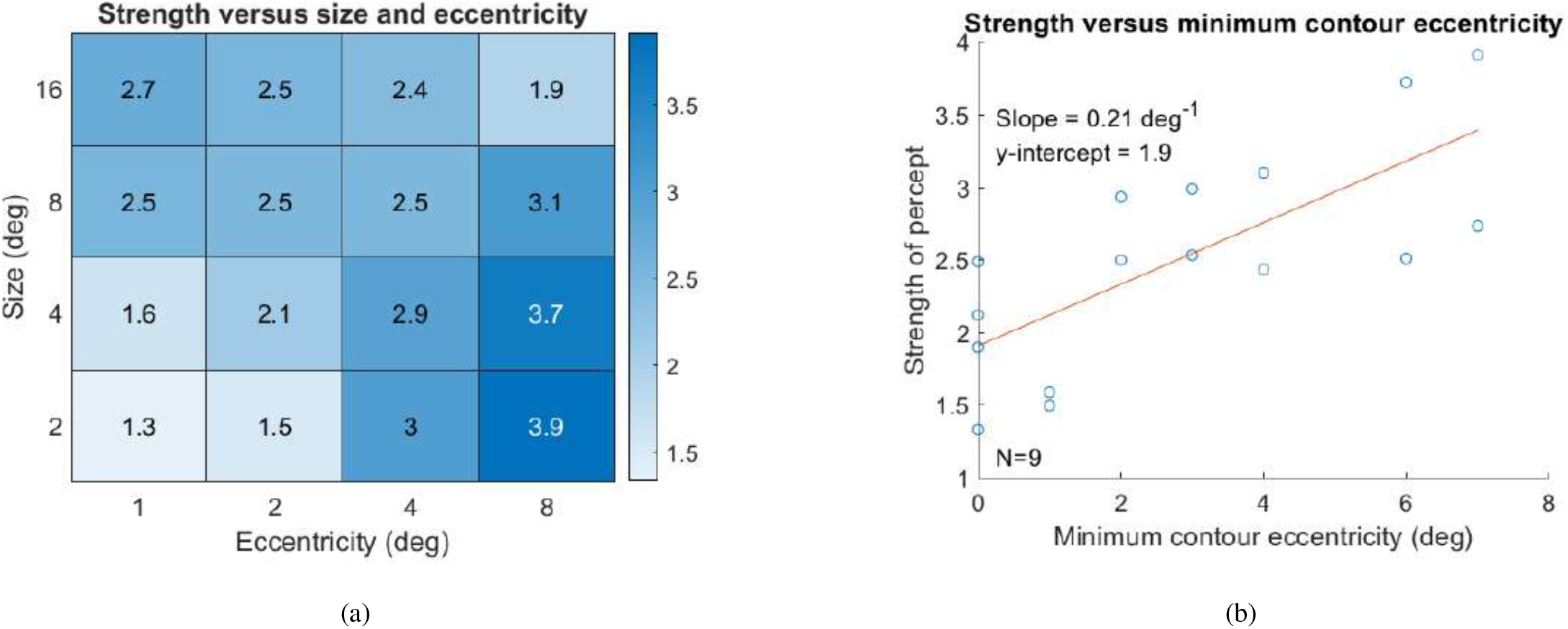
(a) A co-relational matrix shows the strength as a function of size and eccentricity (b) Strength versus minimum contour eccentricity: The strength of the percept strengthens as we move away from the fovea. Strength values are means across all participants.

Large objects at small eccentricities and small objects at large eccentricities produce large effects. Small objects at small eccentricities and large objects at large eccentricities produce smaller effects. This is demonstrated in the co-relation matrix in Figure 7 (a). The minimum contour eccentricity measures the eccentricity of the closest part of the contour from the fovea center or the fixation cross. Figure 7(b) shows that the further the contour is from the fovea, the stronger the effect.

Using fitlme, we probed (a) if the users influence the strength of the percept (b) if size, as a fixed effect, influences the strength of the percept (c) if eccentricity, as a fixed effect, influences the strength of the percept (d) if both size and eccentricity, influences the strength of the percept (e) if both size, eccentricity and participant as a random effect influences the strength of the percept. We find the following: (a) Users do have an influence on the strength reported. A linear model with strength as the dependent variable and users as the independent variable shows users effect the strength (F =111.9, p *<* 0.05). A one-way Anova on the strength reported by the users shows that users have a large effect (*η* = 0.2516). (b) Size effects the strength reported (*η* = 1.1836e-04). (c) Eccentricity effects the strength reported (*η*=3.6048e-145) (d) Both size and eccentricity effects the size reported (*η*=0.0175). (e) Size, eccentricity and users all effect the strength reported (*η* = 0.26). Size and eccentricity together offer a better explanation of the strength being reported than either effect individually.

To address the concern that the blocks showed possible variation, we tested if block was an effect for the polygon order being reported and found that this was not the case (p *<* 0.05, large negative loglikelihood −454.06).

### Size

The major effect here is stimuli size. Smaller circular stimuli generally show smaller order polygons. Bigger circular stimuli show higher order polygons. This is an interesting feature that indicates that rather than creating large edges between vertices, smaller, more numerous edges are created between numerous vertices.

### Eccentricity

Eccentricity has a modest effect on the polygon order reported. Proximity to the fovea shows higher order polygons. This is a smaller effect. See Figure 8(a),(b)

**Figure 8:**
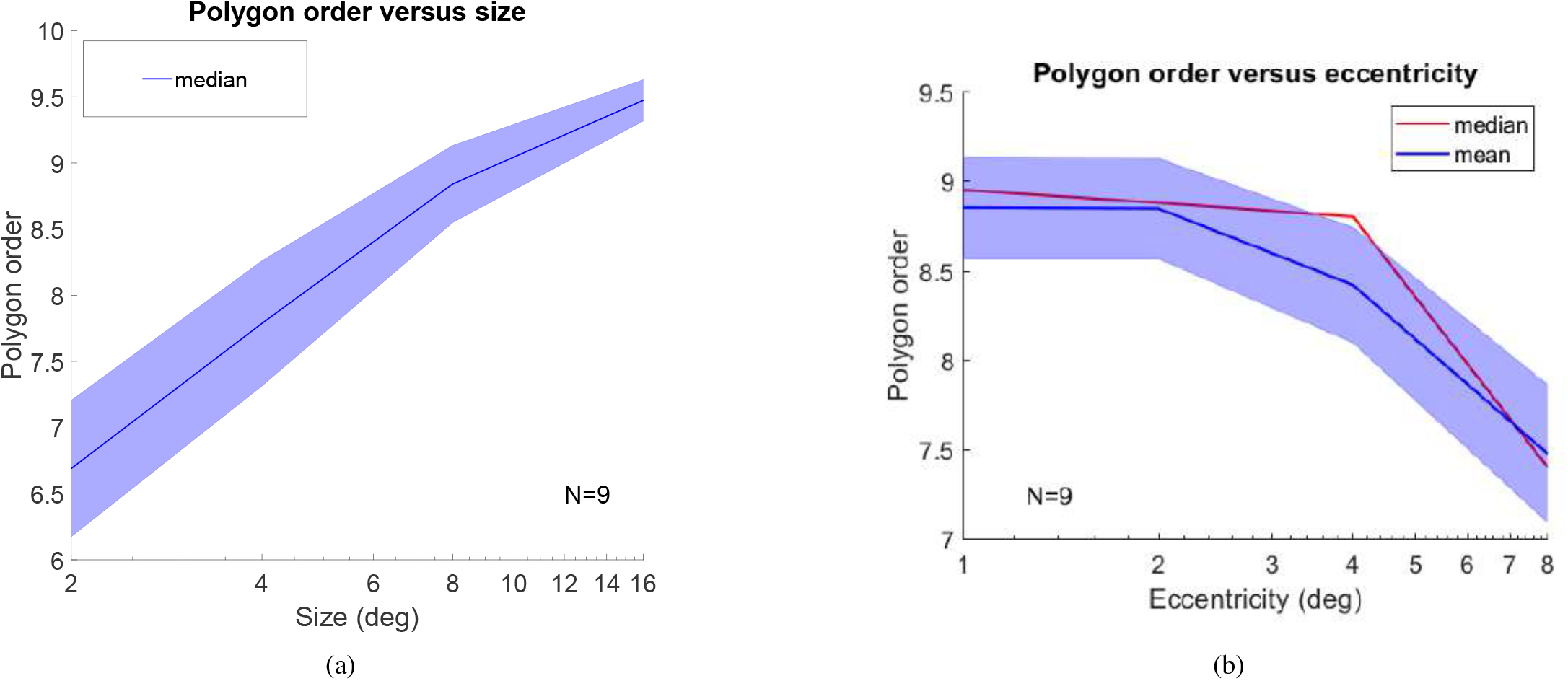
(a) Polygon order versus size: Shaded error bar reflect the standard error of mean around the mean polygon order. Larger order polygons were reported as the diameter of the stimuli increased. (b) Polygon order versus eccentricity: Shaded error bar reflect the standard error of mean around the mean polygon order. In general, observers report lower order polygons at higher eccentricities,larger order polygons at middle eccentricities, followed by slightly lower order polygons at smaller eccentricities. Eccentricity has a modest effect on polygon order. All values reported are means across all participants. The median and mean are shown for eccentricity only since they are not as closely overlapping and show slight user differences.

**Figure 9:**
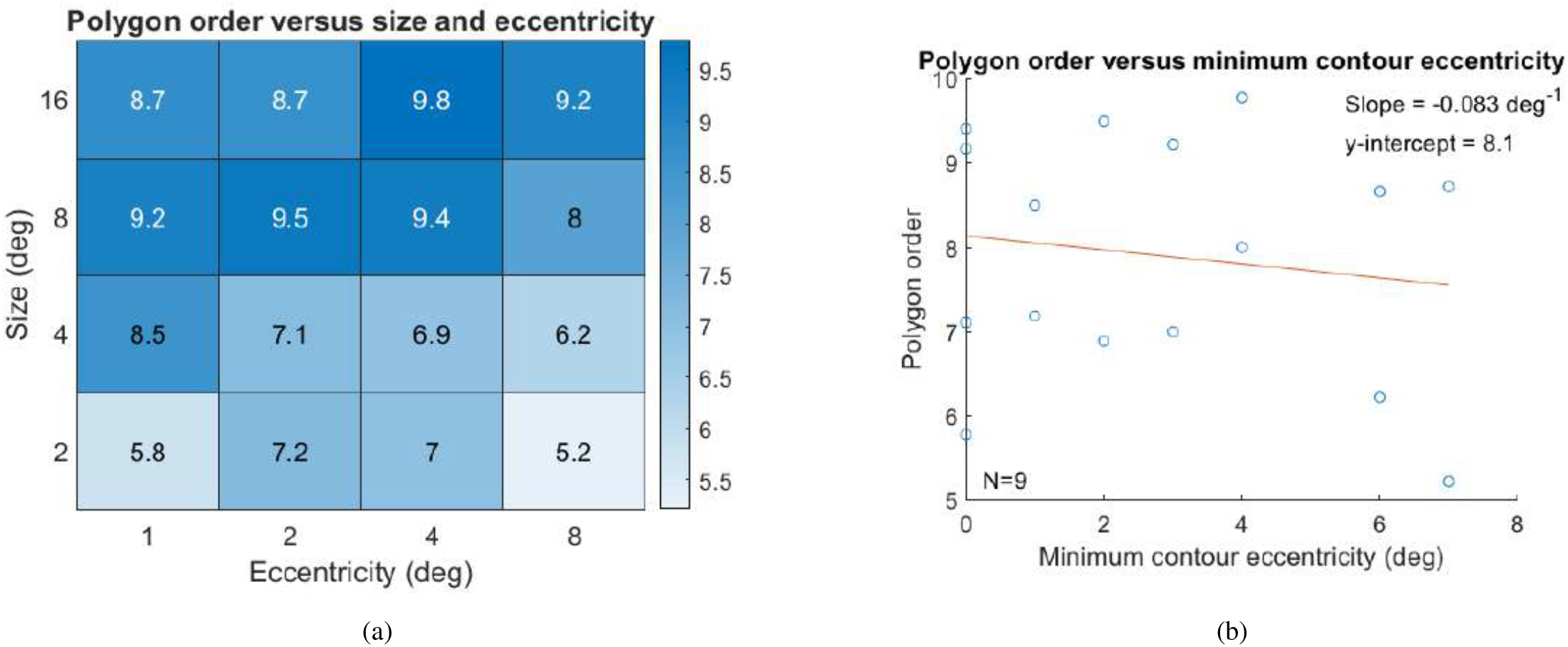
(a) Co-relation matrix showing polygon order (mean) versus size and eccentricity. (b) Polygon order versus minimum contour eccentricity. On average, the minimum contour eccentricity from the fovea only modestly effects the polygon order. Means are over all participants.

### Orientation

We compare observer responses to a uniform distribution of 15 angle increments starting from 0 to the first axis of symmetry for the polygon in question. For most polygon orders, a uniform distribution is observed across all orientations of the polygon (Figure 10. The triangle showed a relatively high KL divergence value compared to other polygons. With the triangle, observers preferred the 45 ^°^ orientation strongly. Figure 10 shows the responses recorded and the orientation selected for the response polygon.

**Figure 10:**
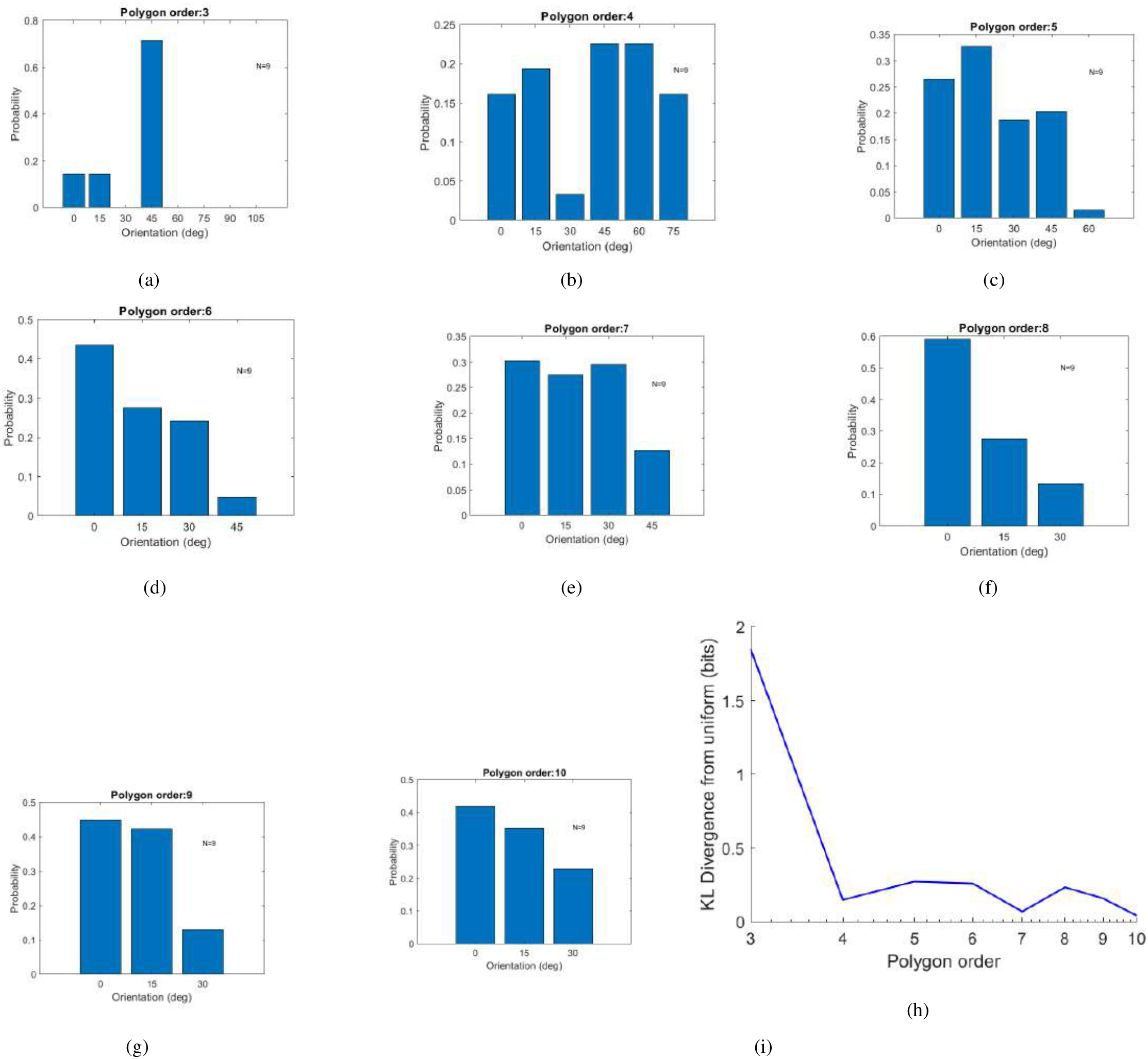
Probability versus orientation : (a-h) For each polygon that was selected by the user, we show the distribution of the orientations selected. The canonical form (0 deg orientation) is when the polygon is centred with its apex at the 12 o’clock position. The responses are computed over all users, shown as a probability of selecting that orientation over all the orientations selected per polygon. In Figure 3(b,c), the first shape in the orientation array is the canonical representation of the triangle and square respectively. (i) KL divergence: KL divergence shows how much the orientation differs from a uniform distribution of orientations. When observers select a triangle as the in-place adaptation polygon, they mostly select the 45 degree orientations. Other shapes show a more uniform spread in the orientations selected.

**Figure 11:**
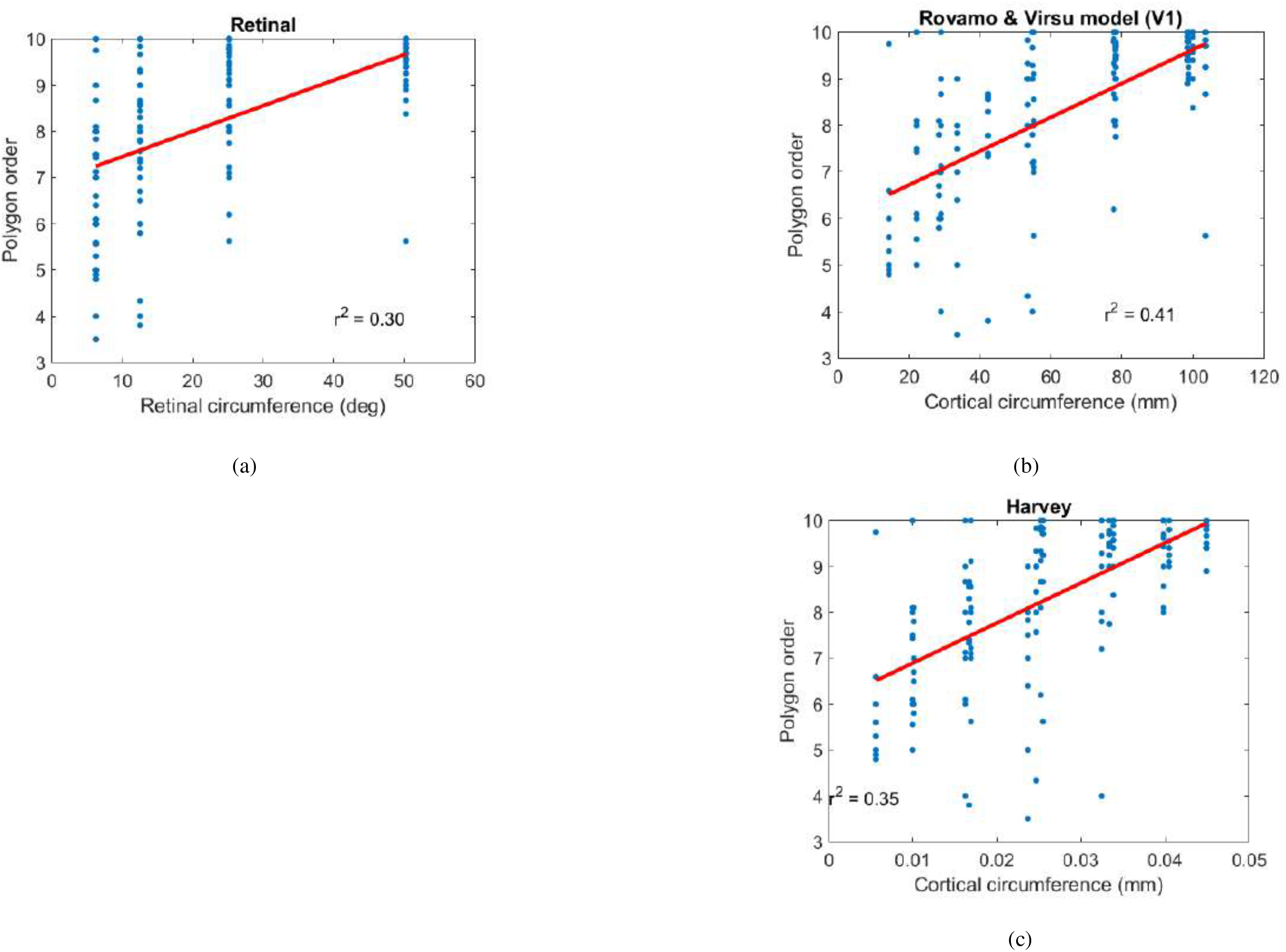
(a) Retinal model: Polygon order regressed over retinal circumference. (b) Rovamo and Virsu cortical model: Polygon order regressed over retinal circumference using Rovamo and Virsu’s cortical magnification factor (CMF) for region V1. (c) Harvey cortical model: Polygon order regressed over cortical circumference using Harvey’s Cortical Magnification Factor for V4.

Preference for contour orientation, where the contour is pointing in a particular angle and direction, is a known phenomenon in mid-level vision, specifically in region V4 (Pasupathy & Connor, 1999a, 2001, 2002). However, the responses recorded do not see any emergent internal polygon angle, after fatiguing of the circular contour, that is angled or oriented as part of its constituent contour shape as shown in (Pasupathy & Connor, 1999a, 2001, 2002). Pasupathy and Connor (Pasupathy & Connor, 1999b) show that there is a neural preference for angles oriented in the 135 ^°^ to 180 ^°^ direction. However, in the tests conducted, we find no preference for any particular orientation across polygons (Figure 10(i)).

### Model that explains the polygon order observed

Using regression, we build an equation for the polygon order as a function of the retinal circumference (Equation 2) and as a function of the cortical circumference (Equation 3).

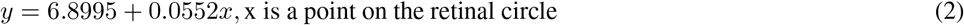

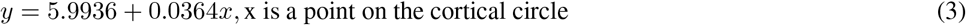

The *R*^2^ value explains how well the model explains the variability in the responses based on the variability in the predictor values. The cortical length’s *R*^2^ reports higher value than the *R*^2^ for the retinal length measurements. We used bootstrapping on the observer’s responses i.e. polygon order and computed the retinal function and cortical function *R*^2^ values. Among the models that we studied - (a) the skeletal model with Rovamo and Virsu cortical magnification factor and (b) the Harvey model - we found *R*^2^ values are the highest with the skeletal model using the Rovamo and Virsu cortical magnification factors. This model, therefore, becomes our model of choice in explaining the type of polygon seen during adaptation. The p-value measures what proportion of times *R*^2^ is bigger for the retinal than the cortical function. The cortical function *R*^2^ of 1 compared favourably over a retinal function *R*^2^ value of 0. See Figure 12. This proves that our brain samples the curve at points along the circle’s circumference which is better explained by a cortical length function.

**Figure 12:**
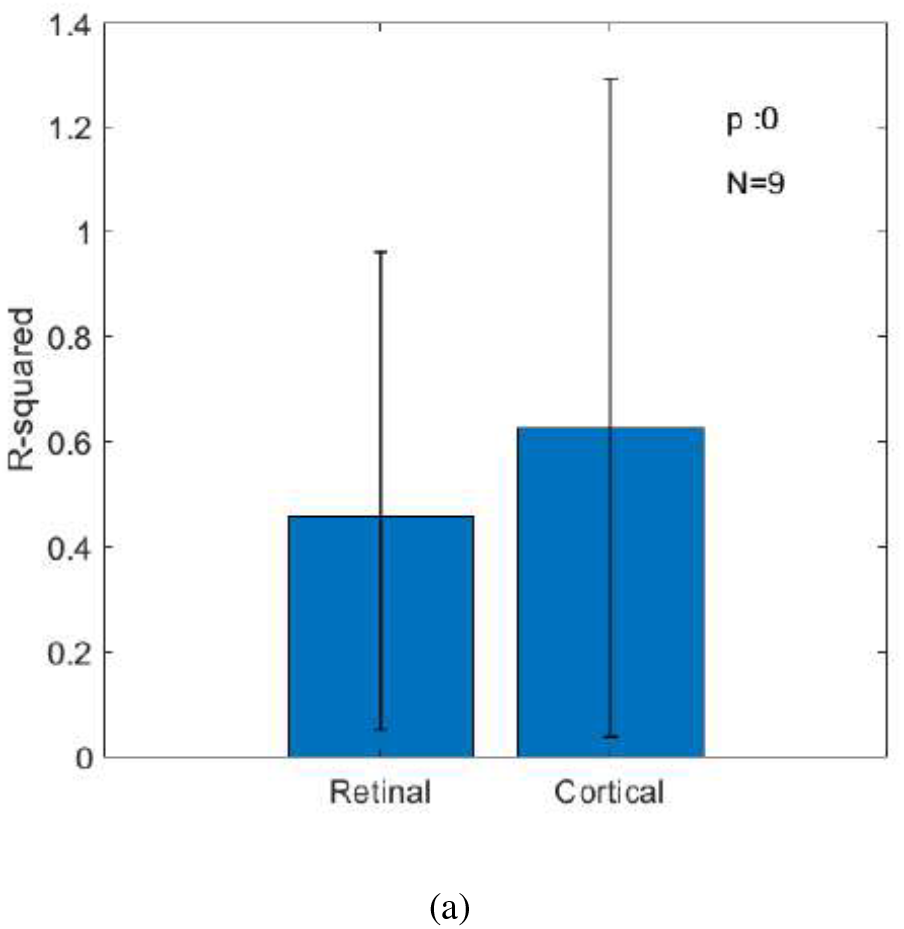
R-squared, from 10000 bootstrapped samples, for retinal circumference function and cortical circumference function. p-value measures by how much *R*^2^ for the retinal circumference function is greater than *R*^2^ for the cortical circumference function. Proportion times retinal is greater than cortical is 0. Proportion times cortical is greater than retinal is 1.

As evident from regression, a better summary of the relationship between the polygon order seen during adaptation is provided by a cortical function than a retinal function in the brain’s region V1 using Rovamo and Virsu’s magnification factors. The cortical magnification factors used from lower level V1 (Rovamo and Virsu model) offers a better fit than the cortical magnification factors used from higher regions from the brain such as V4 (Harvey model).

## Discussion

We make an important observation about scale invariance in this paper. As polygon order changes with changing size, it shows that scale invariance, a distinct property of the visual system, is not obeyed, with respect to curvature processing. Scale invariance is a feature of the higher order visual system. A low-level account explains an opponency between localized curvature-tuned neurons in early visual cortex (Das & Gilbert, 1995, 1997), while the higher-level account assumes an opponency between neurons in higher visual areas coding whole shapes (El-Shamayleh & Pasupathy, 2016a). Because the high-level account that would predict invariance to size and position fails, the low-level account that predicts that the order of the perceived polygon will increase with size and decrease with the eccentricity of the stimulus, due to cortical magnification, becomes a plausible theory.

Several studies have attempted to understand how the curve is mechanistically coded. Curvature built by a neural population is widely favoured because curvature appearance is explained by neurons tuned to different curvatures (Ben-Shahar & Zucker, 2004; Gheorghiu & Kingdom, 2017). Neural populations often define the shapes of afterimages, because it is often the aggregated inhibition responses, collectively that influence what is seen as an afterimage (Gheorghiu & Kingdom, 2017). In our experiment, we show that the unique formation of regular polygons during the adaptation process is a manifestation of the early cortical computation in region V1. The simplest mechanistic model that describes the polygon is defined by the equation given by the regression equation,

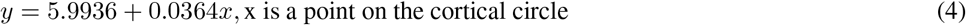

The equations confirms that the area that best explains the appearance of polygons is region V1 of the cortex.

The participants were asked to reason about strength and shape (in terms of polygon order). The strength confirmed percepts were strong enough to report reliably.

The central visual field responds to a higher change in orientation, i.e. higher curvature, since the central receptive fields are smaller in all dimensions than the peripheral receptive fields (D. Hubel & Wiesel, 1974). The fact that we see higher order polygons closer to the fovea than the periphery aligns with this theory.

Rotation invariance is observed with respect to normalized curvature (El-Shamayleh & Pasupathy, 2016b). Normalized curvature is described as the rate of change in tangent angle per unit of angular length. Absolute curvature is described as the rate of change in tangent angle with respect to contour length. When the shape scales up or down, the normalized curvature remains the same while the absolute curvature changes.A large set of neurons, coding for normalized curvature, maintain selectivity for the shape across size changes whereas a smaller proportion show a shift in selectivity for shape with changing size. Neurons firing for normalized curvature describe orientation that scales with size whereas neurons firing for absolute curvature describe orientation at points described by linear distances. The higher mid-level V4 neurons, with larger receptive fields, shows preference for certain angle (Pasupathy & Connor, 2001, 2002; El-Shamayleh & Pasupathy, 2016a). However, there is no cohesive selection for orientation across stimulus orders.

Future work can explore the stimuli as gradually increasing arcs of the circle to see how much of the local curvature i.e. arc circumference is needed to code the polygon seen. El-Shamayleh et. al in (El-Shamayleh & Pasupathy, 2016b) show that scale invariance can be explained by neurons that respond to normalized curvature instead of absolute curvature. This raises interesting questions about the lower limits of normalized curvature coding by the population neurons and the upper limit of local curvature coding by the population neurons when polygon order changes with size. These limits are exposed during adaptation when the neurons are fatigued in upcoming future work.

## Acknowledgements

The authors would like to thank Bob Ho for setting up the apparatus and Nicholas Baker for graciously reading this manuscript and offering valuable editorial suggestions.

